# Pupil-linked arousal does not differ between ‘white’, ‘pink’ and ‘brown’ noises

**DOI:** 10.1101/2025.06.11.659177

**Authors:** Mercede Erfanian, Maria Chait, Jian Kang

## Abstract

‘Coloured’ noises, such as white, pink, and brown noise, have gained attention in popular media as potential tools for enhancing memory consolidation, sleep quality, attentional focus, and more. These terms refer to distinct spectral slopes, which give rise to perceptually different noise stimuli. Although empirical research on their effects remains limited, a prevailing hypothesis suggests that their influence may be mediated by differential effects on arousal. In this study, we investigated this hypothesis using pupillometry, a physiological marker closely linked to arousal. Participants (*N* = 31) listened to three types of noise (white, pink, and brown), each presented for 10 sec, while their pupil diameter was recorded. The results showed no significant modulation of pupil size across noise conditions. These findings suggest that, despite widespread claims about the distinct arousing or calming properties of coloured noises, they do not differentially affect sustained pupil-linked arousal.

## Introduction

In recent years, predominantly driven by reports in popular media ^1, 2, 3^, there has been growing public interest in ‘coloured’ noises, such as white, pink and brown noise, as aids for relaxation and sleep. These sounds are widely promoted for their soothing properties among both neurotypical individuals and those with neurodiversity (Attarha et al., 2018; Ebben et al., 2021; Nigg et al., 2024; Ong et al., 2016; Riedy et al., 2021; Zhou et al., 2012). However, empirical research confirming these effects or explaining their underlying mechanisms remains limited.

The term ‘colour’ in this context refers to the spectral properties of noise signals that arise from stochastic processes (Doob, 1942). Specifically, these noises differ in their spectral slope, leading to distinct perceptual experiences (Kitapci et al., 2021). White noise features equal power across all frequency bands (1/*f* ^*0*^), while pink noise (1/*f* ^*1*^) exhibits decreasing power with increasing frequency, and brown noise (1/*f* ^*2*^) emphasizes low-frequency components (De Coensel et al., 2003; Gilden et al., 1995; Voss, 1992; Voss, 1975; Yang et al., 2015) (**Figure 1**).

**Figure 1.**
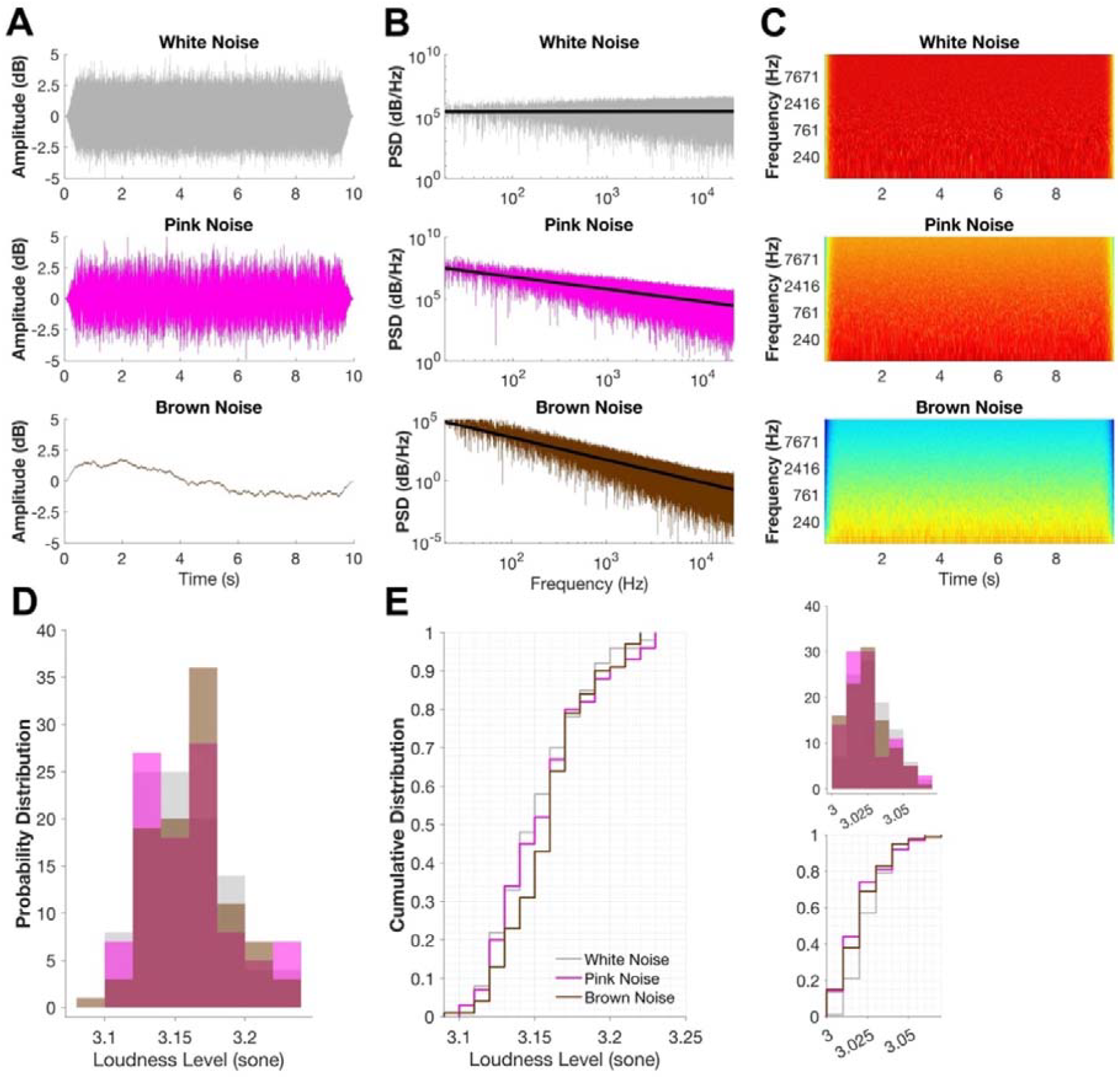
A: Example waveforms of white, pink, and brown noise. B: Power spectral density (PSD) in log-log scale with spectral diagonal slopes reflecting characteristic 1/f^α^ spectral scaling. C: The spectra for the three stimulus conditions with logarithmically spaced frequency axes. D: Probability density distribution, and E: cumulative distribution of short-term loudness; insets display the corresponding long-term distributions.

Despite ongoing debate about their efficacy, coloured noises are widely used in applied contexts, including masking unwanted environmental sounds (Ebben et al., 2021; Loewen et al., 1992; Schäffer et al., 2018), managing tinnitus (Bauer et al., 2011; Coles et al., 1984; Krick et al., 2015; Lai et al., 2023), and improving sleep quality and duration by inducing low-frequency brain oscillations (Ebben et al., 2021; Lee et al., 2024; Lu et al., 2020; Zhou et al., 2012). Furthermore, some studies suggest that exposure to coloured noise can enhance cognitive functions, including memory, by modulating beta and gamma oscillations, key frequencies involved in working memory (Hatayama et al., 2021; Lundqvist et al., 2016). Coloured noise may also enhance learning by strengthening connectivity between midbrain regions and the Superior Temporal Sulcus (STS), critical for attention modulation, via dopaminergic neuromodulation (Hatayama et al., 2021; Lu et al., 2020; Pinardi et al., 2023; Rausch et al., 2014; Shin et al., 2013). However, preferences for different colour noise appear highly subjective, and the physiological basis for these preferences, presumably linked to their spectral characteristics, remains poorly understood.

The heightened arousal often associated with white noise may be due to its dense high-frequency content which is often perceived as louder (Henoch et al., 1999). Indeed, high-pass filtering increases arousal (Buono et al., 2021), whereas low-pass filtering reduces it (Schorradt et al., 2018).

Due to their 1/*f* spectral properties, these noise types also differ in their degree of spectral stochasticity. White noise is completely uncorrelated and unpredictable; pink noise exhibits moderate predictability; and brown noise is highly dependent on past states, resulting in a structured and predictable signal (De Coensel et al., 2003; R. Voss, 1992; R. F. Voss, 1992; Voss et al., 1978; Voss, 1975; Yang et al., 2015). A substantial body of literature, typically focused on the time domain, has examined how predictability, even in rapid temporal patterns, influences perception. Predictable inputs are generally easier for the brain to process (Milne et al., 2021; Rohenkohl et al., 2012) and to suppress (Andreou et al., 2011; Makov et al., 2020; Southwell et al., 2017), suggesting that they impose a lower demand on attentional resources. If similar mechanisms apply to spectral predictability, this could help explain the perceptual and cognitive effects commonly associated with exposure to colored noise.

In this study, we use the well-established link between pupil dilation and arousal to test the hypothesis that the perceptual effects of coloured noise are mediated by differences in arousal. Non-luminance-mediated pupil dilation, or the Pupillary Dilation Response (PDR), is (at least partly) associated with noradrenaline release from the locus coeruleus (LC), a brain stem nucleus involved in regulating arousal and vigilance (Aston-Jones et al., 2005; Joshi et al., 2016). Prior studies have linked increased pupil dilation with high-frequency auditory content (Wetzel et al., 2016) and unpredictability, which is thought to reflect increased processing demands (Friedman et al., 1973; Huviyetli & Chait, 2025; Milne et al., 2021; Qiyuan et al., 1985). Specifically, Milne et al. (2021) and Huviyetli & Chait (2025) have shown that rapid, regularly repeating, predictable auditory patterns elicit sustained reduced pupil responses compared to matched random sequences, suggesting that predictable inputs release processing resources, resulting in lower arousal.

If the 1/*f* ^*o*^ slope of white noise, which combines high-frequency content and low predictability, does indeed elevate arousal, we would predict greater pupil dilation in response to white noise relative to pink and brown noise, and potentially a further graded difference between the latter two. To test this, we conducted a lab-based experiment in which participants listened to matched-duration (10 s) white, pink, and brown noise stimuli while their pupil size was continuously recorded. All procedures were aligned with prior studies in which arousal-related effects on sustained pupil dilation were observed (Huviyetli & Chait, 2025; Milne et al., 2021) in order to maximize the likelihood of detecting an effect, should one exist. To control for loudness as a potential confound, given that sounds with broader frequency spectra are typically judged as louder, all stimuli were equalized for perceived loudness. This approach allows us to isolate the effect of spectral ‘colour’ (1/*f* slope) on arousal-related pupil dynamics.

## Materials and Methods

### Participants

Thirty-eight paid participants were invited to take part in the experiment. The data of seven participants were excluded from the analysis (see Participants exclusion criteria). The data of thirty-one participants are included in the analysis (22 females, mean age = 26.52 ± 4.53 years, range = 19-35). All participants reported that they had no known otological or neurologic disorders/conditions and had normal or corrected-to-normal vision, with Sphere (SPH) prescriptions not exceeding 3.5. The study was approved by the research ethics committee of University College London and written informed consent was obtained from all participants.

Previous work using a similar analysis of tonic pupil responses ((Milne et al., 2021); *N* = 20) demonstrated robust effects with a smaller sample size. Therefore, the current sample size is considered sufficient to detect meaningful effects.

### Participant exclusion criteria

Participants were removed from the final analysis if more than 50% of their trials were classified as invalid (‘bad trials’, see below). Based on this criterion, five participants were excluded due to excessive trial-level artifacts. Two additional participants were excluded for non-compliance with task instructions. Non-compliance instances included a lack of attention to study instructions, such as failing to gaze at the fixation cross, maintaining stillness during the experiment, and responding appropriately to incidental silent gaps (see below) by pressing the designated key.

### Stimuli

Stimuli were 10 s noises (white, pink, and brown) with 500 ms cosine on- and offset ramps. Signals were generated by setting a PSD of 1/*f*^*α*^ over its entire frequency range, using a sampling frequency of 44100 Hz. The inverse frequency power was *α* = 0 for white, *α* = 1 for pink, and *α* = 2 for brown noises as suggested by (Zhivomirov, 2018). Overall, 100 instances of each condition were generated. To control for perceived loudness, the loudness of each stimulus was computed and normalized using the model proposed by Moore-Glasberg ((Moore et al., 2016); ISO 532-2 (2017)), implemented via the 32-bit & 64-bit Loudness PROGRAMS. This procedure was iteratively applied until the stimuli fell within a predefined loudness target range (long-term loudness: 3.00 –3.06 sone; short-term loudness: 3.09 – 3.23 sone). The selected range was arbitrary and chosen to achieve a reasonable degree of uniformity in loudness across stimuli, without reference to specific perceptual thresholds.

**Figure1D and E** demonstrate the probability density and cumulative distributions of the short-term loudness and long-term loudness (insets) of coloured noises in sone (*N* = 300) obtained from the generated brown noises (mean short-term = 3.16 ± 0.03 and mean long-term = 3.02 ± 0.01), pink noises (mean short-term = 3.15 ± 0.03 and mean long-term = 3.02 ± 0.01), and white noises (mean short-term = 3.15 ± 0.02 and mean long-term = 3.02 ± 0.01). A series of Kolmogorov-Smirnov (KS) tests indicated no significant difference between distributions for long-term (brown and pink D = 0.05, *p* = 0.98, brown and white D = 0.1, *p* = 0.68, and pink and white D = 0.14, *p* = 0.26), and short-term (brown and pink D = 0.14, *p* = 0.26, brown and white D = 0.17, *p* = 0.09, pink and white D = 0.06, *p* = 0.98). The results suggest that all coloured noises were successfully normalized to the same range of loudness.

### Experiment

Participants were instructed to sit with their heads placed on a chinrest in front of a monitor (24-inch BENQ XL2420T with a resolution of 1920 × 1080 pixels and a refresh rate of 60 Hz), in a dimly lit and acoustically shielded room (IAC triple-walled sound-attenuating booth). Noises (auditory stimuli) were presented diotically with Sennheiser HD558 headphones (Sennheiser) via a Roland DUO-CAPTURE EX USB Audio Interface (Roland Ltd). Stimuli were presented at a comfortable level (∼ 65 dB A) which was determined during piloting, and all participants confirmed it was appropriate. Stimulus presentation and response recording were controlled with Psychtoolbox (Psychophysics Toolbox version 3; (Brainard, 1997) on MATLAB (version 2019a) (MATLAB, 2019)).

#### Behavioural task

Attention to the auditory stimuli was quantified as performance on an incidental task (as in (Huviyetli & Chait, 2025; Milne et al., 2021)). Participants were instructed to monitor the presented noises for brief silent gaps. This ensured participants remained broadly attentive without imposing a high cognitive load. The typical threshold for detecting gaps in noise is approximately 2-5 ms (Samelli et al., 2008); we used gaps of 50 ms to create highly salient and easily detectable targets. The silent gaps were placed at a random position between 2 s post-stimulus onset to 2 s pre-stimulus offset between ∼17% to 25% of the noise sequences.

#### Main experiment

In the main experiment, participants were presented with a black fixation cross on a grey background. They were instructed to fixate on the cross while monitoring for the silent gaps, and to respond by pressing the ‘space bar’ as quickly as possible when a gap was detected. Visual feedback was given at the end of each trial. At the end of each block, feedback regarding overall performance was provided, indicating the percentage of correct and incorrect responses. The experiment comprised of six blocks, each lasting approximately 8 min, along with a practice block. Each block included 15 trials without embedded silent gaps (five trials per condition: ‘brown’, ‘pink’, and ‘white’) and between three to five trials per block containing silent gaps. This resulted in a total of 108 to 120 trials per participant, including 18 to 30 trials with embedded silent gaps and 90 trials without silent gaps (30 trials per condition). The stimulus set for each participant was selected randomly without replacement from a pool of pre-generated stimuli (a total of 100 ‘main’ stimuli and 25 stimuli with a gap for each ‘colour’).

Inter-trial intervals were jittered between 5000 and 7000 ms. Stimuli were all presented in a random order. The experiment lasted ∼75 mins. Breaks were provided between blocks for 3 to 5 mins.

### Gap detection task analysis

Sensitivity scores (d’) were calculated by using the hit and FA rate d’ = z (HR) – z (FA). A keypress would be considered a hit if it occurred <1.5 s following the silent gap. When HR or FA were at the ceiling (or value of 1 and value of 0 respectively, results in undefined d’), the *loglinear* correction was applied (Hautus, 1995). The overall task performance was high with the HR close to ceiling (brown = 99.6%, pink = 97.98%, and white = 97.95%), while FA base rate was low (brown = 3.86%, pink = 2.72%, white = 3.19%). Since d’ data were not normally distributed, a Friedman test was conducted to compare the performance across conditions. Reaction times (RT) were recorded from each ‘hit’ and were similarly analysed using the Friedman test **(Figure 2A, B, C, D)**.

**Figure 2.**
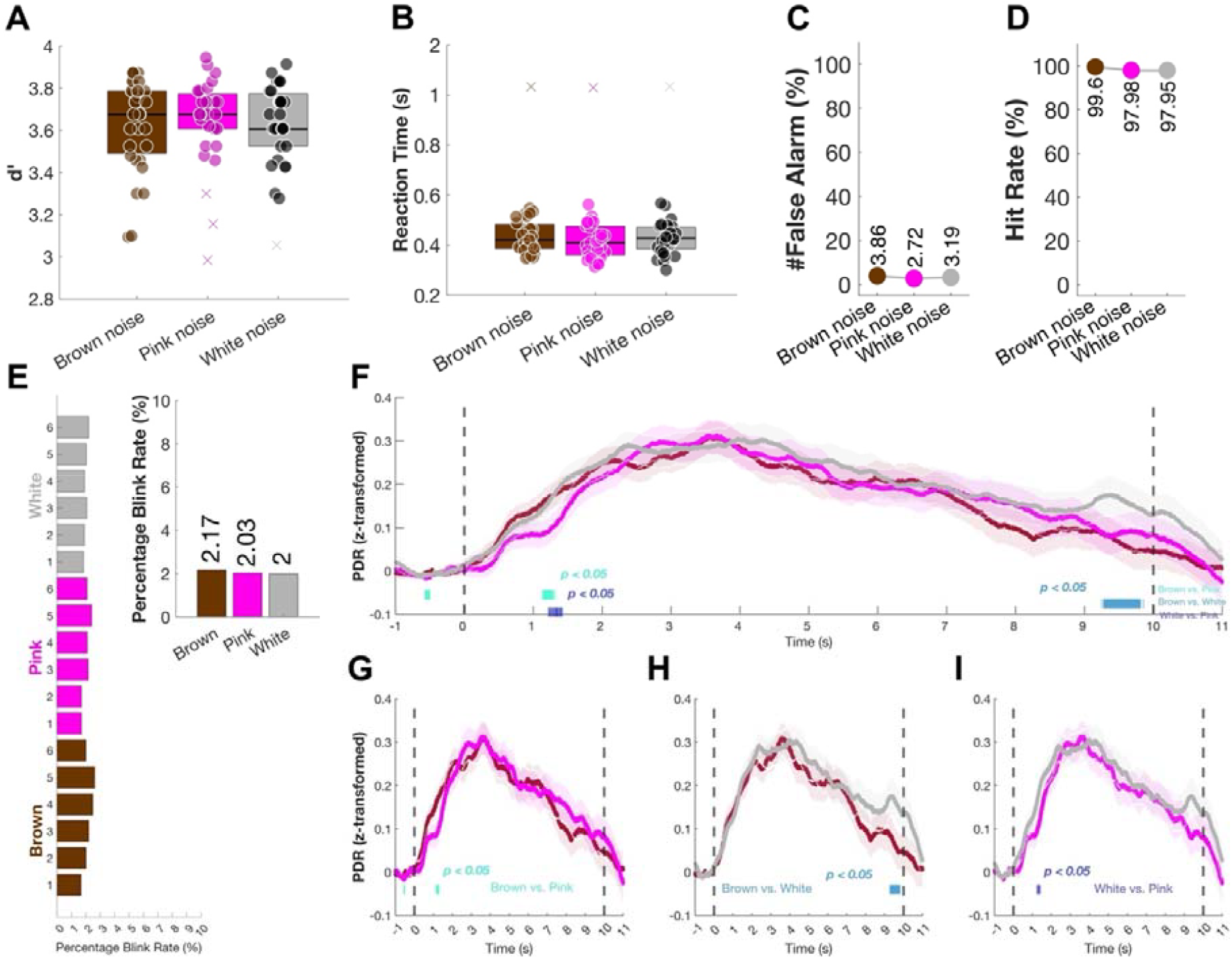
A – D: Gap detection performance across noise conditions. In A and B, circles denote individual participant data. E shows blink rates for each block (1–6) across the three noise conditions (e.g., White), with percentage blink rate on the x-axis. The inset displays the overall mean blink rate for each condition. F-I overlayed pupil dilation responses to ‘brown’, ‘pink’, and ‘white’ noise stimuli are shown. Shaded areas represent ±1 SEM. Horizontal bars (light to dark blue) indicate significant differences (p<⍰0.05). The analysis was also conducted at p< 0.01, no significant intervals were found.

### Pupil diameter measurement

An infrared eye-tracking camera (Eyelink 1000 Desktop Mount, SR Research Ltd.) was situated at a horizontal distance of 65⍰cm away from the subject. The standard 5-point calibration procedure for the Eyelink system was conducted prior to each experimental block. During the experiment, the eye-tracker continuously tracked gaze position and pupil diameter focusing binocularly at a sampling rate of 1000⍰Hz. Participants were instructed to blink normally and rest their eyes during intervals. Additional instruction was provided to the participants with excessive blinking or those who failed to fixate appropriately. Prior to each trial, the eye tracker automatically checked that the participants’ eyes were open and fixating; the trial would not start if this was not confirmed.

#### Baseline pupil measure

Prior to the main experiment, participants underwent a pupil reactivity assessment to evaluate basic autonomic responsivity. This included measuring steady-state pupil diameter across gradual luminance shifts (white to black) including transient pupillary responses to sudden luminance increase (pupillary light reflex), luminance decrease (pupillary dark reflex), and brief auditory stimuli (harmonic tones). These procedures served to confirm normal pupil dynamics and screen for atypical responsivity (Wang et al., 2018). No participants showed atypical pupillary responses during the screening procedures.

#### Pre-processing of pupillometry data

Trials with silent gaps and trials where the participants made a FA were excluded from further analysis.

The left eye data were analysed. For the time series analysis, the PDR to the noises were epoched from 1 s prior to the noise onset to 11 s post-stimulus onset (a total window width of 12 s). Blinks or partial eye closure were detected and recovered by using shape-preserving piecewise cubic interpolation. The average blink rate (proportion of excluded samples due to eye closure) was ∼2% which slightly increased toward the end of the experiment **(Figure 2E)**.

In order to allow comparison across subjects, data for each subject in each block were Z normalized by calculating the mean and standard deviation across all baseline samples (1 s pre-onset interval) in that block. Subsequently, for each participant, pupillary dilation was time-domain averaged across all epochs of each condition (‘white’, ‘pink’ and ‘brown’) to generate a single time series per condition.

We adopted a conservative approach for detecting ‘bad trial’s (e.g., outliers). Outlier trials, defined as those exceeding ±2 SD from the condition-specific mean, were identified and removed separately within each experimental condition to preserve condition-related variability. Trials with over 50% missing data (e.g., due to blinks or partial eye closure) were also excluded. However, no trials exceeded this threshold in any condition.

#### Time series statistical analysis

A non-parametric bootstrap-based analysis (Efron et al., 1994) was conducted to examine time-domain differences between conditions. For each participant, the difference in time series between each pair of conditions was calculated and the time series were subjected to bootstrap resampling (1000 iterations with replacement). Significant differences at each time point were determined based on whether the proportion of bootstrap iterations falling below or above zero exceeded 99% (*p*< 0.01).

## Results

### Gap detection task results

Sensitivity to the silent gaps was analysed by using d’ **(Figure 2A)**. Overall, as expected, performance was at ceiling, with no differences between conditions in d’ χ2(2) = 1.19, *p* = 0.55 or RT χ2(2) = 2.39, *p* = 0.3.

### Pupil size as a measure of tonic arousal does not vary with noise type

**Figure 2F** plots the average PDR as a function of time, relative to the pre-onset baseline (1 s). All three noises elicited almost similar PDR patterns. Slightly after the stimulus onset (*t* ∼0.5), the pupillary dilation rapidly increased and peaked within 3 s. Pupil diameter then entered a prototypical phase of slow reduction. This is similar to the pattern observed in other studies using long stimuli (Huviyetli & Chait, 2025; Milne et al., 2021; Zhao et al., 2019). The statistical analysis indicated no difference between conditions using *p*< 0.01. While brief periods of significance were observed at *p*< 0.05, their short duration suggests they likely reflect noise rather than a reliable effect of pupil-linked arousal which we expect to be sustained in nature.

## Discussion

We investigated the hypothesis that pupil-linked arousal differs across coloured noise types, in an effort to understand the mechanisms underlying their reported efficacy in promoting relaxation. Our results revealed no measurable differences in pupil size between white, pink, and brown noise conditions, suggesting that the perceived effects of these noises are not mediated by general arousal levels as assessed via pupillometry.

Previous studies using similar paradigms have demonstrated that bottom-up driven arousal-related changes can indeed be detected in pupil size. For instance, Milne et al. (2021) and Huviyetli & Chait (2025) reported that sustained pupil diameter was significantly smaller in response to predictable auditory patterns compared to random ones. This effect emerged approximately 3 s after stimulus onset and persisted throughout the stimulus duration (9 s in those studies). These findings were interpreted to suggest that the detection of regularity in the auditory system, even when it is not behaviourally relevant, leads to reduced global arousal, as reflected in reduced pupil dilation. Based on the hypothesis that white noise is associated with increased arousal, we expected it to elicit a larger sustained pupil diameter than pink or brown noise, which are commonly linked with more calming effect. However, as shown in **Figure 5**, this effect was not observed in our data.

One possibility is that the null result is due to the relatively short stimulus duration used in our experiment (10 s), compared to the extended durations (often hours) employed in real-world contexts where these noises are used to influence arousal (Kucukoglu et al., 2016; Suzuki et al., 1991; Zhou et al., 2012). Nevertheless, the spectral differences between these noises are theoretically detectable by the auditory system within milliseconds to seconds, certainly faster than the timescale required for regularity effects previously reported (Milne et al., 2021). Therefore, if these spectral properties were influencing arousal directly, we would expect to observe corresponding differences in pupil size within the time frame used here. The absence of such an effect may suggest that the perceptual or physiological influence of coloured noise unfolds via slower or different mechanisms or may depend on interactions with other factors such as loudness, which was carefully controlled in this study.

Another plausible interpretation of our findings lies in the widespread and roughly equal popularity of all coloured noise types across both neurotypical and neurodivergent populations. Although formal incidence data is lacking, online platforms report substantial numbers of downloads, clicks, and streams for white, pink, and brown noise alike. This broad usage may indicate that: (a) the three noise types are similarly positioned along affective dimensions such as pleasantness and arousal; (b) no single coloured noise is consistently preferred over others in perceptual terms; (c) individual preferences are less about the acoustic properties themselves and more about how well the noise satisfies specific functional needs such as masking specific unwanted environmental sounds (Ebben et al., 2021; Loewen et al., 1992).

In summary, the present study suggests that, under controlled laboratory conditions and using a 10 s stimulation period, there is no measurable difference in pupil-linked arousal across different noise types. This finding constrains existing hypotheses about the mechanisms underlying the efficacy of these noises and rules out a simple baseline explanation that the three noise categories differ in terms of automatic, bottom-up arousal responses. Instead, it shifts the focus toward alternative pathways through which these sounds may exert their reported soothing effects, such as cognitive, contextual, or top-down mechanisms.

## Notes

**Declaration of Conflicting Interests** The authors declared no potential conflicts of interest with respect to the research, authorship, and/or publication of this article.

**CRediT Statement** ME conceptualized the study and developed the methodology with input from MC and JK. ME conducted data collection, performed pre-processing, formal analysis, and curated the data under the supervision of MC. ME drafted the original manuscript, with editing conducted collaboratively and technical feedback by MC. MC and JK supervised the project. Additionally, MC and JK oversaw project administration and supported funding acquisition efforts.

https://osf.io/uz5fa/

### Competing Interest Statement

The authors have declared no competing interest.

## References

Andreou, L. V., Kashino, M., & Chait, M. (2011). The role of temporal regularity in auditory segregation. Hear Res, 280(1-2), 228–235. 10.1016/j.heares.2011.06.001

Aston-Jones, G., & Cohen, J. D. (2005). An integrative theory of locus coeruleus-norepinephrine function: adaptive gain and optimal performance. Annu Rev Neurosci, 28, 403–450. 10.1146/annurev.neuro.28.061604.135709

Attarha, M., Bigelow, J., & Merzenich, M. M. (2018). Unintended consequences of white noise therapy for tinnitus—otolaryngology’s cobra effect: a review. JAMA Otolaryngology–Head & Neck Surgery, 144(10), 938–943. 10.1001/jamaoto.2018.1856

Bauer, C. A., & Brozoski, T. J. (2011). Effect of tinnitus retraining therapy on the loudness and annoyance of tinnitus: a controlled trial. Ear and Hearing, 32(2), 145–155. 10.1097/AUD.0b013e3181f5374f

Brainard, D. H. (1997). The Psychophysics Toolbox. Spat Vis, 10(4), 433–436. <https://www.n>cbi.nlm.nih.gov/pubmed/9176952

Buono, G. H., Crukley, J., Hornsby, B. W., & Picou, E. M. (2021). Loss of high-or low-frequency audibility can partially explain effects of hearing loss on emotional responses to non-speech sounds. Hearing research, 401, 108153. 10.1016/j.heares.2020.108153

Coles, R., Baskill, J. L., & Sheldrake, J. B. (1984). Measurement and management of tinnitus*: Part I. Measurement. The Journal of Laryngology & Otology, 98(12), 1171–1176. 10.1017/s0022215100148248

De Coensel, B., Botteldooren, D., & De Muer, T. (2003). 1/f noise in rural and urban soundscapes. Acta acustica united with acustica, 89(2), 287–295.

Doob, J. L. (1942). What is a stochastic process? The American Mathematical Monthly, 49(10), 648–653. 10.1080/00029890.1942.11991300

Ebben, M. R., Yan, P., & Krieger, A. C. (2021). The effects of white noise on sleep and duration in individuals living in a high noise environment in New York City. Sleep medicine, 83, 256–259. 10.1016/j.sleep.2021.03.031

Efron, B., & Tibshirani, R. J. (1994). An introduction to the bootstrap. Chapman and Hall/CRC. 10.1201/9780429246593

Friedman, D., Hakerem, G., Sutton, S., & Fleiss, J. L. (1973). Effect of stimulus uncertainty on the pupillary dilation response and the vertex evoked potential. Electroencephalography and clinical neurophysiology, 34(5), 475–484.

Gilden, D. L., Thornton, T., & Mallon, M. W. (1995). 1/f noise in human cognition. Science, 267(5205), 1837–1839. 10.1126/science.7892611

Hatayama, A., Matsubara, A., Nakashima, S., & Nishifuji, S. (2021). Effect of pink noise on EEG and memory performance in memory task. 2021 IEEE 10th Global Conference on Consumer Electronics (GCCE),

Hautus, M. J. (1995). Corrections for extreme proportions and their biasing effects on estimated values of d1. Behavior research methods, instruments, & computers, 27, 46–51.

Henoch, M. A., & Chesky, K. (1999). Ear canal resonance as a risk factor in music-induced hearing loss. Medical Problems of Performing Artists, 14(3), 103–106.

Huviyetli, M., & Chait, M. (2025). The Interplay of Bottom-Up Arousal and Attentional Capture during Auditory Scene Analysis: Evidence from Ocular Dynamics. BioRxiv, 2025.2004. 2025.650619.

Joshi, S., Li, Y., Kalwani, R. M., & Gold, J. I. (2016). Relationships between pupil diameter and neuronal activity in the locus coeruleus, colliculi, and cingulate cortex. Neuron, 89(1), 221–234. 10.1016/j.neuron.2015.11.028

Kitapci, K., & Akbay, S. (2021). Audio-visual interactions and the influence of colour on noise annoyance evaluations. Acoustics Australia, 49(2), 293–304. 10.1007/s40857-021-00220-x

Krick, C. M., Grapp, M., Daneshvar-Talebi, J., Reith, W., Plinkert, P. K., & Bolay, H. V. (2015). Cortical reorganization in recent-onset tinnitus patients by the Heidelberg Model of Music Therapy. Frontiers in Neuroscience, 9, 49. 10.3389/fnins.2015.00049

Kucukoglu, S., Aytekin, A., Celebioglu, A., Celebi, A., Caner, I., & Maden, R. (2016). Effect of White Noise in Relieving Vaccination Pain in Premature Infants. Pain Manag Nurs, 17(6), 392–400. 10.1016/j.pmn.2016.08.006

Lai, H., Wang, G., Zheng, Z., Gao, M., Li, S., & Wu, S. (2023). Pink noise: a potential sound therapy for tinnitus. American Journal of Translational Research, 15(11), 6621.

Lee, H.-A.-N., Lee, W.-J., Kim, S.-U., Kim, H., Ahn, M., Kim, J.-H., Kim, D.-W., Yun, C.-H., & Hwang, H.-J. (2024). Effect of Dynamic Binaural Beats on Sleep Quality: A Proof-of-Concept Study with Questionnaire and Biosignals. Sleep, zsae097. 10.1093/sleep/zsae097

Loewen, L. J., & Suedfeld, P. (1992). Cognitive and arousal effects of masking office noise. Environment and Behavior, 24(3), 381–395. 10.1177/0013916592243006

Lu, S. Y., Huang, Y. H., & Lin, K. Y. (2020). Spectral content (colour) of noise exposure affects work efficiency. Noise Health, 22(104), 19–27. 10.4103/nah.NAH_61_18

Lundqvist, M., Rose, J., Herman, P., Brincat, S. L., Buschman, T. J., & Miller, E. K. (2016). Gamma and Beta Bursts Underlie Working Memory. Neuron, 90(1), 152–164. 10.1016/j.neuron.2016.02.028

Makov, S., & Zion Golumbic, E. (2020). Irrelevant Predictions: Distractor Rhythmicity Modulates Neural Encoding in Auditory Cortex. Cereb Cortex, 30(11), 5792–5805. 10.1093/cercor/bhaa153

MATLAB. (2019). Release, Statistics Toolbox, The MathWorks, Inc., Natick, Massachusetts, United States. In: ed.

Milne, A. E., Zhao, S., Tampakaki, C., Bury, G., & Chait, M. (2021). Sustained Pupil Responses Are Modulated by Predictability of Auditory Sequences. J Neurosci, 41(28), 6116–6127. 10.1523/JNEUROSCI.2879-20.2021

Moore, B. C., Glasberg, B. R., Varathanathan, A., & Schlittenlacher, J. (2016). A loudness model for time-varying sounds incorporating binaural inhibition. Trends in hearing, 20, 2331216516682698. 10.1177/2331216516682698

Nigg, J. T., Bruton, A., Kozlowski, M. B., Johnstone, J. M., & Karalunas, S. L. (2024). Systematic Review and Meta-Analysis: Do White Noise and Pink Noise Help With Attention in Attention-Deficit/Hyperactivity Disorder? Journal of the American Academy of Child & Adolescent Psychiatry. 10.1016/j.jaac.2023.12.014

Ong, J. L., Lo, J. C., Chee, N. I., Santostasi, G., Paller, K. A., Zee, P. C., & Chee, M. W. (2016). Effects of phase-locked acoustic stimulation during a nap on EEG spectra and declarative memory consolidation. Sleep medicine, 20, 88–97. 10.1016/j.sleep.2015.10.016

Pinardi, M., Schuler, A. L., Arcara, G., Ferreri, F., Marinazzo, D., Di Pino, G., & Pellegrino, G. (2023). Reduced connectivity of primary auditory and motor cortices during exposure to auditory white noise. Neurosci Lett, 804, 137212. 10.1016/j.neulet.2023.137212

Qiyuan, J., Richer, F., Wagoner, B. L., & Beatty, J. (1985). The pupil and stimulus probability. Psychophysiology, 22(5), 530–534. 10.1111/j.1469-8986.1985.tb01645.x

Rausch, V. H., Bauch, E. M., & Bunzeck, N. (2014). White noise improves learning by modulating activity in dopaminergic midbrain regions and right superior temporal sulcus. J Cogn Neurosci, 26(7), 1469–1480. 10.1162/jocn_a_00537

Riedy, S. M., Smith, M. G., Rocha, S., & Basner, M. (2021). Noise as a sleep aid: a systematic review. Sleep Medicine Reviews, 55, 101385. 10.1016/j.smrv.2020.101385

Rohenkohl, G., Cravo, A. M., Wyart, V., & Nobre, A. C. (2012). Temporal expectation improves the quality of sensory information. J Neurosci, 32(24), 8424–8428. 10.1523/JNEUROSCI.0804-12.2012

Samelli, A. G., & Schochat, E. (2008). The gaps-in-noise test: gap detection thresholds in normal-hearing young adults. Int J Audiol, 47(5), 238–245. 10.1080/14992020801908244

Schäffer, B., Pieren, R., Schlittmeier, S. J., & Brink, M. (2018). Effects of different spectral shapes and amplitude modulation of broadband noise on annoyance reactions in a controlled listening experiment. International journal of environmental research and public health, 15(5), 1029. 10.3390/ijerph15051029

Schorradt, M., Castillo, S., & Cunningham, D. W. (2018). The semantic space for emotional speech and the influence of different methods for prosody isolation on its perception. Proceedings of the 15th ACM Symposium on Applied Perception,

Shin, S.-K., & Shim, J.-Y. (2013). Relationship between effects of pink noise on brain wave concentration index by individual characteristics and multiple intelligence. Science of Emotion and Sensibility, 16(4), 481–492. 10.1016/j.jtbi.2012.04.006

Southwell, R., Baumann, A., Gal, C., Barascud, N., Friston, K., & Chait, M. (2017). Is predictability salient? A study of attentional capture by auditory patterns. Philosophical Transactions of the Royal Society B: Biological Sciences, 372(1714), 20160105. 10.1098/rstb.2016.0105

Standard, I. (2017). 532-2: 2017; Acoustics—Methods for Calculating Loudness—Part 2: Moore-Glasberg Method. International Organization for Standardization: Geneva, Switzerland.

Suzuki, S., Kawada, T., Ogawa, M., & Aoki, S. (1991). Sleep deepening effect of steady pink noise. Journal of sound and vibration, 151(3), 407–414. 10.1016/0022-460X(91)90537-T

Voss, R. (1992). 1/f noise and fractals in economic time series. In Fractal Geometry and Computer Graphics (pp. 45–52). Springer.

Voss, R. F. (1992). Evolution of long-range fractal correlations and 1/f noise in DNA base sequences. Physical review letters, 68(25), 3805.

Voss, R. F., & Clarke, J. (1978). ‘‘1/f noise’’in music: Music from 1/f noise. The Journal of the Acoustical Society of America, 63(1), 258–263. 10.1121/1.381721

Voss, R. P. (1975). I/F NOISE”“IN MUSIC and SPEECH.

Wang, C. A., Tworzyanski, L., Huang, J., & Munoz, D. P. (2018). Response anisocoria in the pupillary light and darkness reflex. Eur J Neurosci, 48(11), 3379–3388. 10.1111/ejn.14195

Wetzel, N., Buttelmann, D., Schieler, A., & Widmann, A. (2016). Infant and adult pupil dilation in response to unexpected sounds. Dev Psychobiol, 58(3), 382–392. 10.1002/dev.21377

Yang, M., De Coensel, B., & Kang, J. (2015). Presence of 1/f noise in the temporal structure of psychoacoustic parameters of natural and urban sounds. J Acoust Soc Am, 138(2), 916–927. 10.1121/1.4927033

Zhao, S., Bury, G., Milne, A., & Chait, M. (2019). Pupillometry as an objective measure of sustained attention in young and older listeners. Trends in hearing, 23, 2331216519887815. 10.1177/2331216519887815

Zhivomirov, H. (2018). A method for colored noise generation. Romanian journal of acoustics and vibration, 15(1), 14–19.

Zhou, J., Liu, D., Li, X., Ma, J., Zhang, J., & Fang, J. (2012). Pink noise: effect on complexity synchronization of brain activity and sleep consolidation. Journal of theoretical biology, 306, 68–72. 10.1016/j.jtbi.2012.04.006

